# Rab3a regulates melanin exocytosis induced by keratinocyte-conditioned medium

**DOI:** 10.1101/2021.05.24.445450

**Authors:** Luís C. Cabaço, Liliana Bento-Lopes, Matilde V. Neto, José S. Ramalho, Miguel C. Seabra, Duarte C. Barral

## Abstract

Skin pigmentation relies on melanin and is crucial for photoprotection against ultraviolet radiation-induced toxicity. Melanin is synthesized and stored in melanosomes, within melanocytes and then transferred to keratinocytes. While the molecular players involved in melanogenesis have been extensively studied, those underlying melanin transfer remain poorly characterized. Previously, our group proposed that coupled exo/phagocytosis is the predominant mechanism of melanin transfer in human skin and showed an essential role for Rab11b and the exocyst tethering complex in this process. Using a fluorescence-based assay, we show here that keratinocyte-conditioned medium (KCM) specifically induces melanin exocytosis from melanocytes. Moreover, we found that Rab3a, but not Rab11b, regulates melanin exocytosis upon KCM stimulation. In fact, melanosomes accumulate in melanocyte dendrites upon KCM stimulation, co-localizing with Rab3a mainly in the vicinity of the plasma membrane. Additionally, Rab3a silencing does not affect melanin transfer in melanocyte/keratinocyte co-cultures, in contrast with Rab11b depletion, indicating that Rab11b regulates non-KCM-stimulated melanin exocytosis. Thus, our results suggest the existence of at least two distinct routes of melanin exocytosis: one controlled by Rab11b and another Rab3a-dependent, stimulated by KCM. Furthermore, these results provide evidence that soluble factors secreted by keratinocytes can control skin pigmentation via induction of melanocyte signaling pathways that promote peripheral transport of melanosomes and a Rab3a-mediated exocytosis mechanism.

## INTRODUCTION

The skin pigmentary system ensures protection against ultraviolet radiation (UVr)-induced damage and hence, the onset of skin cancer (Narayanan *et al*., 2010; Bino *et al*., 2018). The photoprotective pigment melanin is synthesized within melanocytes and packaged inside specialized membrane-bound lysosome-related organelles (LROs), termed melanosomes (Marks and Seabra, 2001; Raposo and Marks, 2002; Hearing, 2005). Melanocytes reside in the basal layer of the epidermis sparsely spread at an approximately 1:40 ratio among keratinocytes, forming epidermal-melanin units (Fitzpatrick and Breathnach, 1963; Bino *et al*., 2018). Fully melanized melanosomes are transported from the perinuclear region of melanocytes to the dendrites and then transferred to keratinocytes, where melanin granules form a supra-nuclear cap that absorbs and scatters UVr, protecting nuclear DNA from damage (Scott, 2003; Park *et al*., 2009). Thus, melanin biogenesis and transfer must be tightly regulated to sustain skin pigmentation and ensure efficient photoprotection. Indeed, exposure to UVr enhances secretion of soluble factors by keratinocytes, such as endothelin-1 and α-melanocyte stimulating hormone (α-MSH), which in turn increase melanin synthesis in melanocytes (Yamaguchi and Hearing, 2010; Taisuke and Hearing, 2011; Hirobe, 2014). α-MSH binds to Melanocortin-1 receptor on the surface of melanocytes, stimulating melanogenesis and subsequently increasing melanin transfer to keratinocytes (Mosca et al., 2020). Despite this feedback between melanogenesis and melanin transfer, none of the identified keratinocyte-derived factors has been shown to specifically stimulate melanin transfer from melanocytes to keratinocytes, in addition to the enhancement of melanin synthesis.

The mechanism of melanin transfer remains elusive. We propose that the predominant mode in human skin is coupled exo/phagocytosis (Tarafder *et al*., 2014; Moreiras *et al*., 2021b) ^(1)^. Before melanin is transferred, melanosomes are transported to melanocyte dendrites, where they are tethered to the cortical actin cytoskeleton via the tripartite complex Rab27a/Melanophilin/Myosin Va (Barral and Seabra, 2004; Hume *et al*., 2007). However, at least in two-dimensional melanocyte/keratinocyte co-cultures, Rab27a is not essential for melanin exocytosis and subsequent transfer to keratinocytes (Tarafder *et al*., 2014). In search for potential regulators, we found that Rab11b and the exocyst tethering complex control melanin exocytosis (Tarafder *et al*., 2014; Moreiras *et al*., 2019).

The secretory Rab protein Rab3a was shown by electron microscopy to localize to mature melanosome membranes, although the functional implications of this localization have not been elucidated (Araki *et al*., 2000; Fukuda, 2008). Rab3a is present in melanosome-containing fractions with several soluble N-ethylmaleimide sensitive fusion attachment receptors (SNAREs), including VAMP-2, SNAP-25, SNAP-23, Syntaxin-4 and α-SNAP (Scott and Zhao, 2001). The co-fractionation of SNAREs and Rab3a from mature melanosomes strongly suggests a role for this protein in melanin exocytosis. Rab3a was also described to regulate the trans-SNARE complex assembly, which promotes vesicle fusion with the plasma membrane (Hong, 2005). Indeed, Rab3a is involved in several vesicular secretory processes, such as neurotransmitter release in neurons, hormone release in endocrine cells and insulin secretion in pancreatic β-cells (Fukuda, 2008). Moreover, Rab3 regulates the exocytosis of conventional lysosomes and several LROs, including large dense-core neuroendocrine vesicles, dense core granules and Weibel-Palade bodies (Raposo, G. *et al*., 2007; Fukuda, 2008; Quevedo *et al*., 2018; Delevoye *et al*., 2019). For instance, a Rab3a-dependent complex was described to control lysosome exocytosis during plasma membrane repair (Encarnação *et al*., 2016) and Rab3a is essential for dense core granule exocytosis from sperm cells after Rab27a-mediated transport (Quevedo *et al*., 2018). Furthermore, Rab27, Rab3a and Rab3d show a cooperative role in the secretion of Weibel-Palade bodies from endothelial cells (Raposo, G. *et al*., 2007; Zografou *et al*., 2012; Delevoye *et al*., 2019). In this study, we used a fluorescence-based assay optimized by us to measure exocytosed melanin with increased sensitivity and found that, in addition to stimulating melanogenesis, keratinocyte-conditioned medium (KCM) induces melanin exocytosis from melanocytes in a Rab3a-dependent and Rab11b-independent manner. Thus, we provide evidence to support a KCM-stimulated melanin exocytosis pathway controlled by Rab3a, distinct from Rab11b-dependent non-KCM-stimulated melanin exocytosis.

## RESULTS

### Keratinocyte-conditioned medium increases melanin exocytosis from melanocytes

The most commonly used assay to quantify exocytosed melanin is based on spectrophotometry. This assay has limited sensitivity, which becomes more obvious when quantifying melanin diluted in culture medium. Melanin does not fluoresce in its native state, but it emits fluorescence when oxidized, for instance by H_2_O_2_ (Kayatz *et al*., 2001; Fernandes *et al*., 2016). Based on this property, Fernandes *et al*. established a fluorescence spectrometry method to quantify intracellular melanin, which served as a base for our adaptations (Fernandes *et al*., 2016). We started by evaluating if the culture medium that remains in the isolated pellet of exocytosed melanin affects the measurements. Therefore, considering that the remaining culture medium is diluted 1:8 after the solubilization of the melanin pellet, we assessed the emission spectra at the excitation wavelength of 470 nm. We analyzed different diluted culture media - RPMI, DMEM and KCM (DMEM medium consumed by confluent XB2 keratinocytes in culture for 3 days) - and used NaOH as the blank. All the media showed similar fluorescence intensity along the emission spectra (Figure S1A). Thus, we can confidently state that the culture medium does not interfere with the quantification of exocytosed melanin. Additionally, we observed that synthetic melanin, diluted culture media and the blank show a common emission peak at 550 nm (Figure S1A). Then, to detect the best signal, we optimized both the final volume of sample and the reading distance. These optimizations improved the method’s sensitivity in 100% and we were able to obtain a significant difference between the fluorescence intensity of the blank and 2 μg/ml of synthetic melanin (Figure S1B). Importantly, 2 μg/ml is the approximate concentration of exocytosed melanin after culturing Melan-ink4a melanocytes for 3 days in RPMI-based growth medium (Figure 1A). Finally, we quantified exocytosed melanin and normalized to either total melanocyte number or total protein concentration (Figure S1C). We concluded that the normalization for total protein can be used as a more practical and unbiased approach, since the fold-change between the two samples analyzed is the same using either method (Figure S1C).

**Figure 1:**
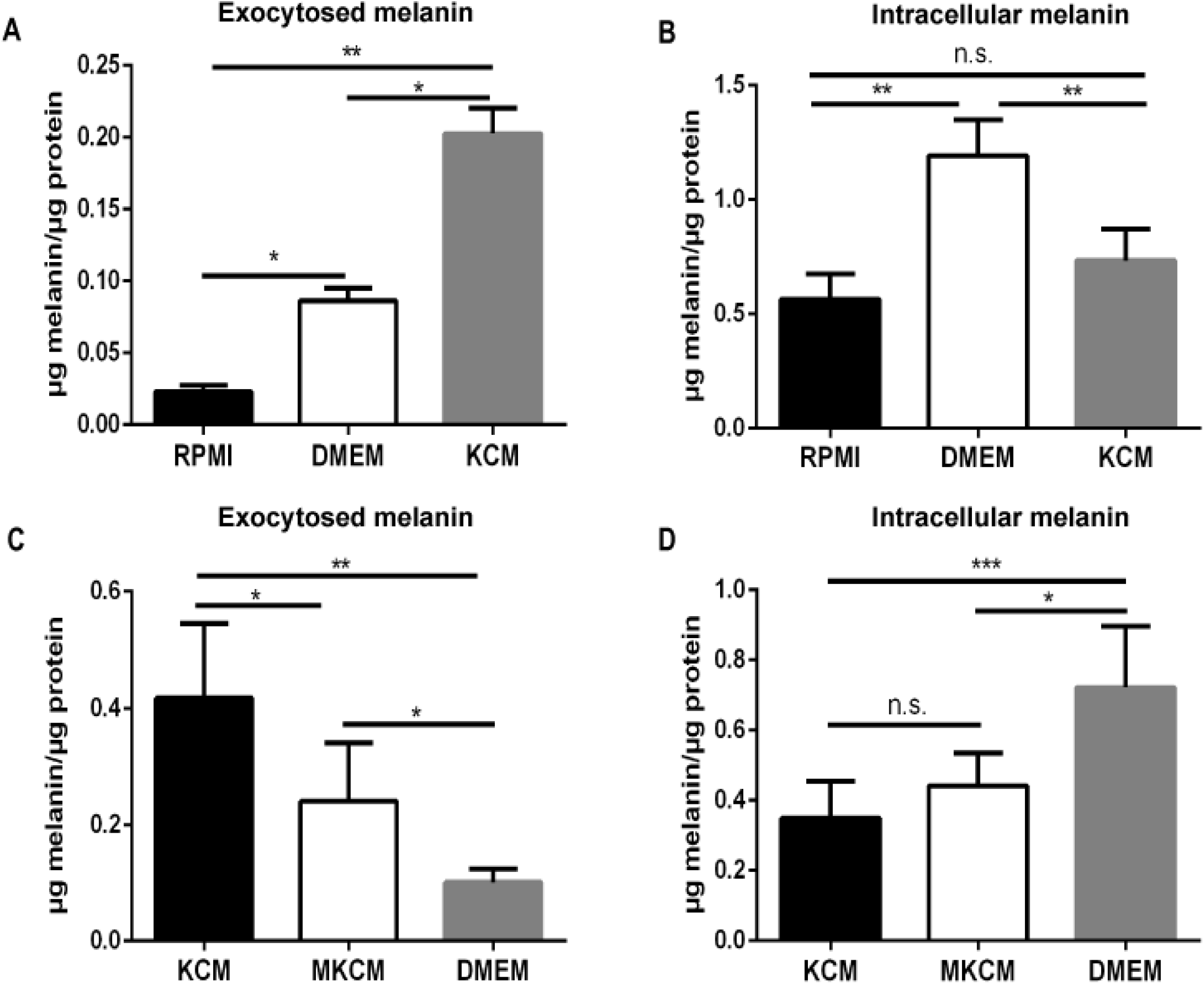
Keratinocyte-conditioned medium stimulates melanin exocytosis from melanocytes. **(A)** Exocytosed and **(B)** intracellular melanin levels of melanocytes cultured with different non-conditioned media (RPMI or DMEM) or keratinocyte-conditioned medium (KCM). **(C)** Exocytosed and **(D)** intracellular melanin levels of melanocytes cultured with DMEM or different DMEM-based conditioned media [KCM or melanocyte/keratinocyte-conditioned medium (MKCM)]. Melanin levels were quantified by fluorescence spectroscopy as described in the Materials and Methods. Results are represented in μg of melanin per μg of total protein. The plots display mean ± SD of at least three independent experiments. One-way ANOVA (n.s. non-significant; * P<0.05; ** P<0.01; *** P<0.0001).

We next aimed to study the stimulation of melanin exocytosis from melanocytes using the newly-established fluorescence-based method. Since keratinocyte-derived soluble factors were shown to increase melanogenesis in melanocytes (Yamaguchi and Hearing, 2010; Taisuke and Hearing, 2011; Hirobe, 2014), we started by evaluating their effect in stimulating melanin exocytosis. To do so, we obtained KCM by culturing XB2 mouse keratinocytes for 3 days and then cultured Melan-ink4a mouse melanocytes with KCM. Both RPMI and DMEM media were used as controls, since the former is the basis of melanocyte growth medium and the latter used to culture keratinocytes. To rule out that the differences observed in exocytosed melanin levels are a consequence of increased melanogenesis, we also quantified intracellular melanin levels.

We first observed that melanocytes cultured in DMEM show a significant increase (3.5-fold) in exocytosed melanin (Figure 1A) and twice the amount of intracellular melanin (Figure 1B), when compared to melanocytes cultured in RPMI. Notably, comparing with RPMI, DMEM has 3.5 times more L-tyrosine, which is the initial substrate of the key melanogenic enzyme tyrosinase (Yamaguchi and Hearing, 2010; Taisuke and Hearing, 2011; Hirobe, 2014). This enhanced intracellular melanin levels in melanocytes cultured with DMEM was also reported by others (Skoniecka *et al*., 2021). Furthermore, we detected a 4-fold increase in melanin exocytosis, when compared with melanocytes cultured with RPMI (Figures 1A and 1B). When melanocytes are cultured in KCM, the levels of exocytosed melanin double (Figures 1A and 1C) and the amount of intracellular melanin decreases by about 50%, compared to melanocytes cultured in DMEM (Figures 1B and 1D). Next, we tested if conditioned medium from melanocyte/keratinocyte co-cultures (MKCM) can stimulate melanin exocytosis in a similar manner to KCM. Surprisingly, we observed that melanocytes cultured with KCM exocytose double the melanin than melanocytes cultured with MKCM (Figure 1C). However, intracellular melanin levels are similar in both cases (Figure 1D). This suggests that keratinocyte-derived factors present in MKCM impact melanin exocytosis in differently than KCM.

### Keratinocyte-conditioned medium stimulates melanin exocytosis in a Rab3a-dependent manner

We next aimed to further dissect the molecular machinery involved in melanin exocytosis from melanocytes, in particular upon KCM stimulation. Previously, we implicated Rab11b in melanin exocytosis from melanocytes cultured with RPMI (Tarafder *et al*., 2014; Moreiras *et al*., 2019). Additionally, Rab3a was suggested to play a role in melanin exocytosis (Araki *et al*., 2000; Scott and Zhao, 2001). Therefore, we analyzed the role of Rab3a and Rab11b in KCM-stimulated melanin exocytosis. For this, we efficiently silenced Rab11b or Rab3a in melanocytes cultured with RPMI, DMEM or KCM (≥60% silencing, Figures S2A and S2B). We also used melanocytes transfected with a non-targeting siRNA (siControl) as the negative control. We found that upon Rab3a depletion, melanin exocytosis is impaired by 50% in melanocytes cultured in KCM, but not in DMEM or RPMI (Figure 2A). In contrast, Rab11b silencing does not affect KCM-induced melanin exocytosis (Figure 2A). Instead, Rab11b silencing significantly reduces melanin exocytosis from melanocytes cultured in RPMI (Figure 2A), confirming our previous reports (Tarafder *et al*., 2014; Moreiras *et al*., 2019). To rule out that the differences observed are due to defects in melanin synthesis, which in turn can impair the amount of exocytosed melanin, we also analyzed the amount of intracellular melanin. We confirmed that the defects in melanin exocytosis cannot be explained by defects in melanin synthesis, since no decrease in intracellular melanin was found in any of the conditions tested (Figure 2B). Instead, we observed a small but significant increase in intracellular melanin levels upon Rab3a depletion in melanocytes cultured with KCM, which could be a consequence of the impairment in melanin exocytosis (Figure 2B). To exclude off-target effects of the siRNA pool used to silence Rab3a, we performed a rescue experiment. For this, we plated wild-type or lentivirus-transduced Melan-ink4a melanocytes overexpressing GFP or GFP-Rab3a and then silenced these cells for Rab3a. Importantly, Rab3a expression levels were assessed by western blot to confirm the silencing and overexpression levels for all the conditions tested (Figures S3A and S3B). We found that by overexpressing human GFP-Rab3a in Rab3a-silenced melanocytes, the amount of exocytosed melanin is restored to the levels observed in siControl melanocytes (Figure 2C). In addition, GFP-Rab3a but not GFP overexpression rescues the augmented intracellular melanin levels caused by Rab3a depletion (Figure 2D). Noteworthy, we found that KCM-stimulated melanocytes overexpressing GFP-Rab3a show double the amount of exocytosed melanin than melanocytes overexpressing GFP (Figure 2E), while no significant differences in intracellular melanin between the two conditions were observed (Figure 2F). This reinforces the conclusion that Rab3a acts as a regulator of KCM-stimulated melanin exocytosis. Altogether, these results suggest the existence of two distinct melanin exocytosis pathways: one regulated by Rab11b and another induced by KCM and regulated by Rab3a.

**Figure 2:**
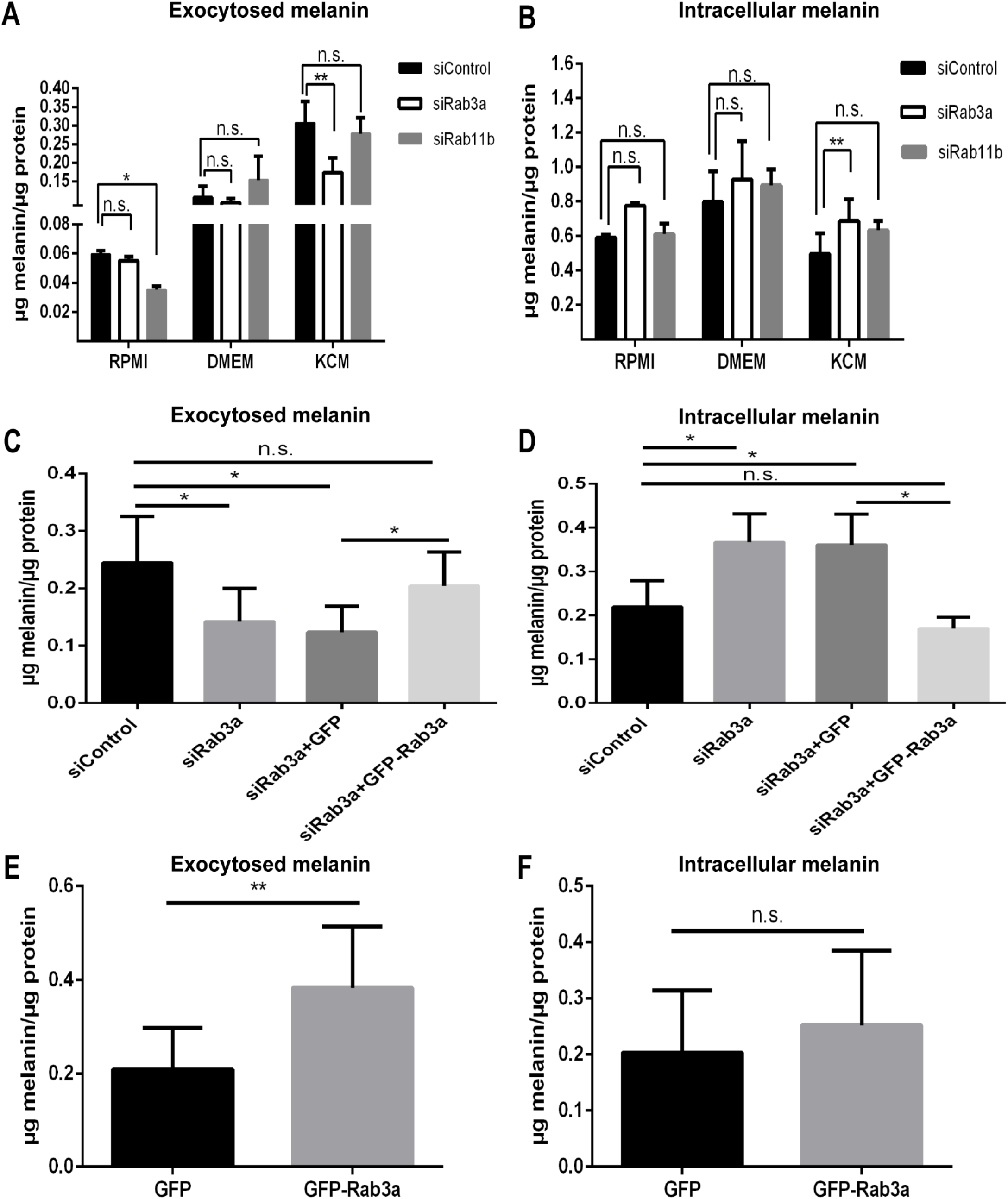
Keratinocyte-conditioned medium induces melanin exocytosis in a Rab3a-dependent and Rab11b-independent manner. **(A)** Exocytosed and **(B)** intracellular melanin levels of melanocytes cultured with RPMI, DMEM or keratinocyte-conditioned medium (KCM) and silenced for Rab3a (siRab3a), Rab11b (siRab11b) or transfected with a non-targeting siRNA (siControl). **(C)** Exocytosed and (**D)** intracellular melanin levels of melanocytes cultured with KCM, overexpressing or not GFP or GFP-Rab3a and silenced for Rab3a or transfected with siControl. **(E)** Exocytosed and **(F)** intracellular melanin levels of melanocytes cultured with KCM and overexpressing GFP or GFP-Rab3a. Melanin amounts (μg) were normalized by total protein (in μg). The plots represent mean ± SD of at least three independent experiments. **(A, B)** Two-way ANOVA; **(C, D)** One-way ANOVA; **(E, F)** Unpaired t-test. n.s. non-significant; *P<0.05; **P<0.01.

### Keratinocyte-conditioned medium promotes melanosome localization to melanocyte dendrites

Since we found that Rab3a regulates KCM-stimulated melanin exocytosis from melanocytes, we next evaluated if Rab3a colocalizes with melanosomes in melanocyte dendrites upon KCM stimulation, to regulate the final steps of melanosome fusion with the plasma membrane. To study this, we overexpressed GFP-Rab3a or GFP as a negative control, in Melan-ink4a melanocytes. The transfected cells were cultured with KCM or DMEM for 24 hours and melanosomes labelled with HMB45 antibody, which recognizes PMEL, a scaffold protein onto which melanin is deposited. Firstly, we confirmed the accumulation of GFP-Rab3a in the Golgi previously described (Zerial and McBride, 2001; Encarnação *et al*., 2016) through co-localization with the coat protein COPI (Figure S4). Then, we evaluated the subcellular localization of melanosomes in melanocytes cultured with DMEM or KCM. In KCM-stimulated melanocytes, 63% of melanosomes are recruited to dendrites and only 37% remain in the cell body (Figures 3A, 3C and 3D), while melanocytes cultured with DMEM show 45% of melanosomes in dendrites and 55% in the cell body (Figures 3B, 3C and 3D). Therefore, the percentage of melanosomes in dendrites increases by about 20% in the presence of KCM, comparing with melanocytes incubated with DMEM (Figures 3C, 4A and 4B). Moreover, KCM increases by about 10% the number of melanosomes positioned close to plasma membrane of dendrites, when compared to DMEM-incubated melanocytes (Figures 4A, 4B, 4D and 4E). Upon overexpression of GFP-Rab3a in KCM-stimulated melanocytes, GFP-Rab3a co-localizes with about 35% of melanosomes in melanocyte dendrites, while only 15% of co-localization is observed in KCM-stimulated melanocytes overexpressing GFP (Figures 4A and 4C). Noteworthy, under KCM stimulation, most GFP-Rab3a co-localizes with melanosomes close to plasma membrane (Figure 4A). These effects appear to be KCM-induced as only about 10% of the melanosomes in the dendrites co-localize with either GFP-Rab3a or GFP in DMEM-incubated melanocytes (Figures 4B and 4C). These results suggest that KCM stimulation induces melanosome transport/tethering to melanocyte dendrites and that Rab3a is recruited to the melanosomes located in the vicinity of the plasma membrane.

**Figure 3:**
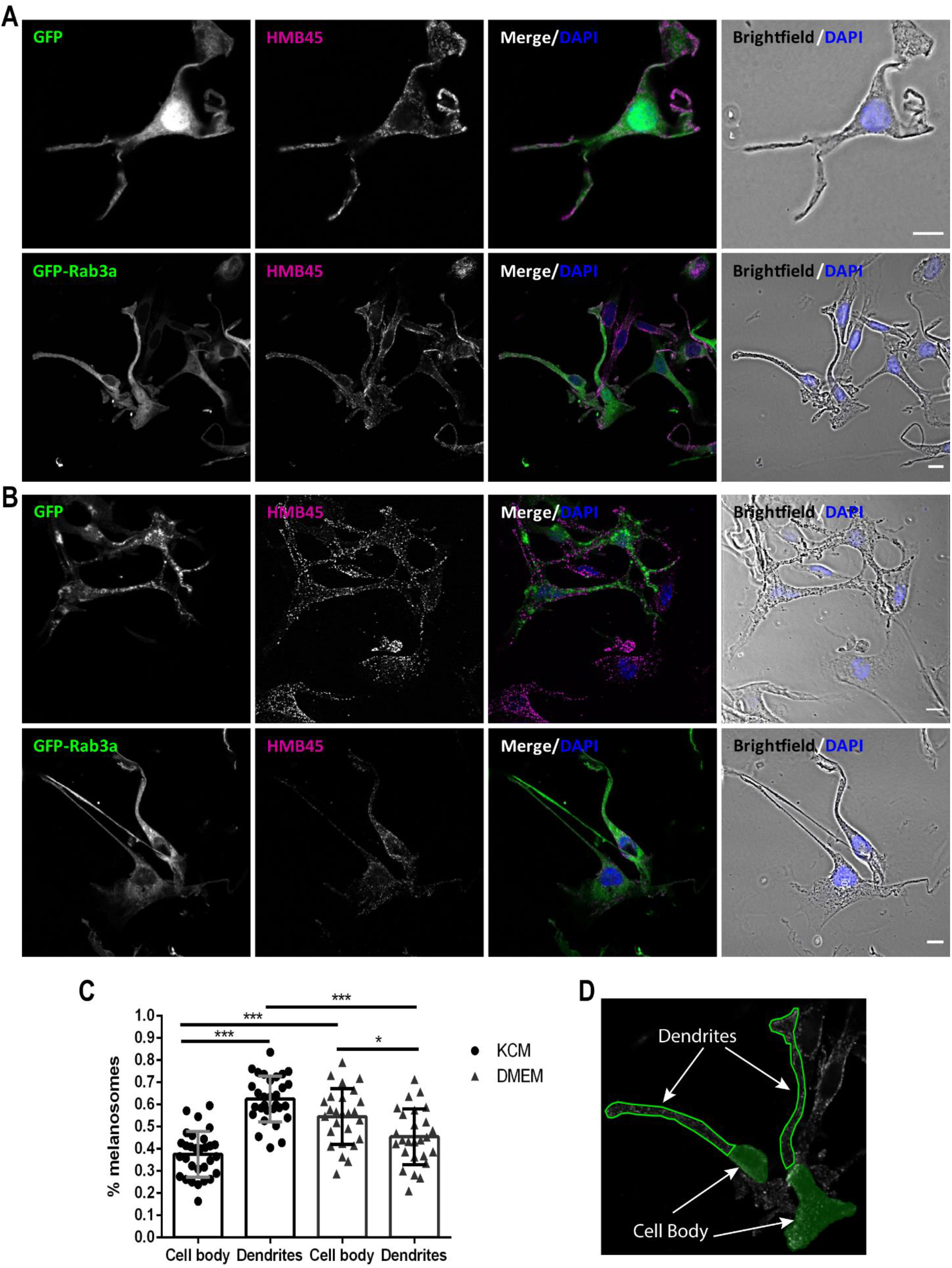
Keratinocyte-conditioned medium increases the number of melanosomes in melanocyte dendrites. Representative confocal images of Melan-ink4a melanocytes cultured with **(A)** keratinocyte-conditioned medium (KCM) or **(B)** DMEM for 24 hours. **(C)** Percentage of melanosomes in the cell body [filled in green, in **(D)**] or dendrites [outlined in green, in **(D)**] of melanocytes cultured with KCM or DMEM. Melanocytes overexpressing GFP or GFP-Rab3a (green) were immunostained to mark melanosomes (pseudocolored in magenta). Nuclei were stained with DAPI (blue). Brightfield shows melanosomes as black dots. Scale bars, 10 µm. The plot displays mean ± SD of three independent experiments. One-way ANOVA (*P<0.05; ***P<0.0001). At least 30 images containing approximately 50 melanocytes in total were analyzed per condition.

**Figure 4:**
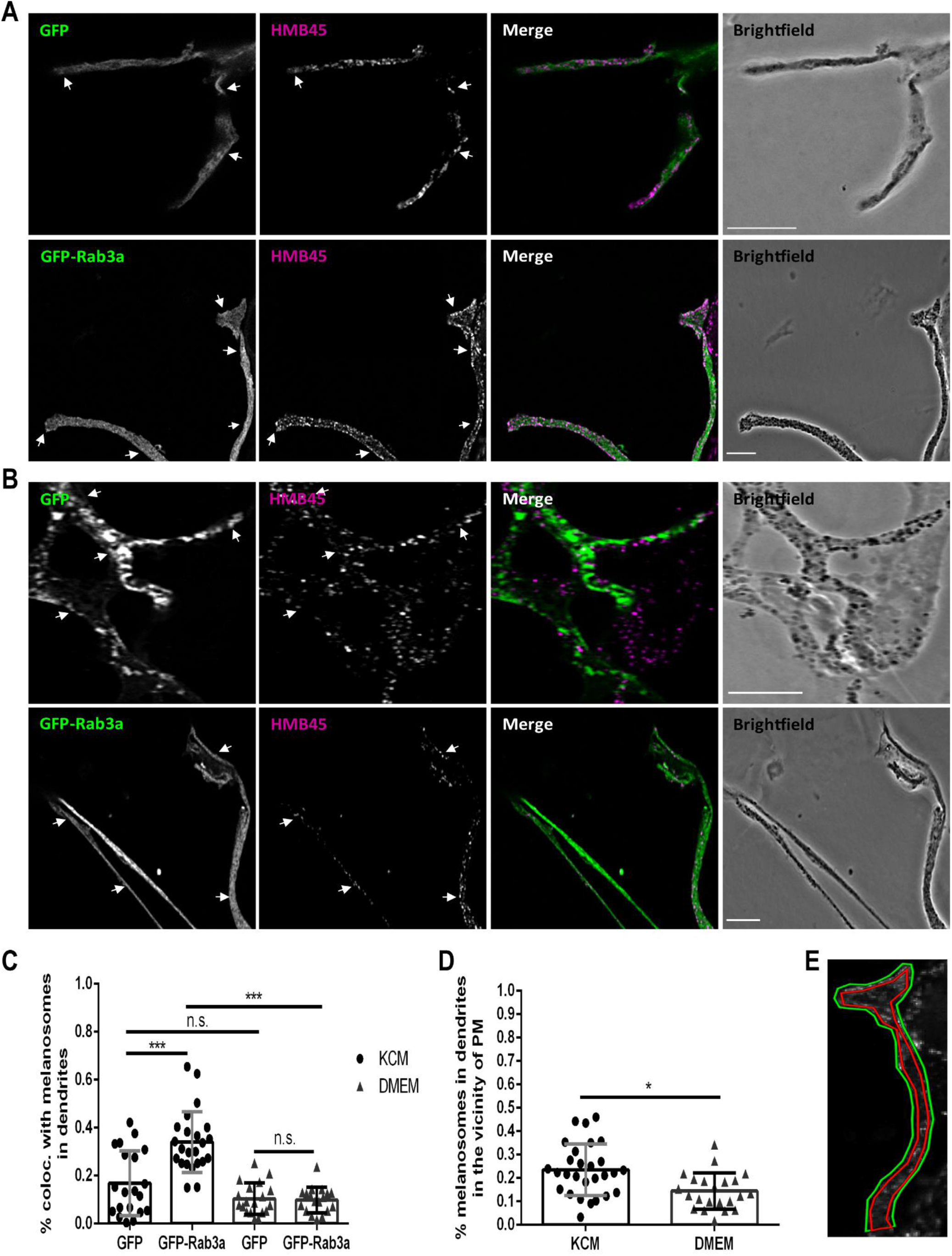
Keratinocyte-conditioned medium promotes co-localization of GFP-Rab3a with melanosomes close to plasma membrane of melanocyte dendrites. Confocal images of Melan-ink4a dendrites cultured with **(A)** keratinocyte-conditioned medium (KCM) or **(B)** DMEM. **(C)** Mander’s coefficient of co-localization (coloc.) between melanosomes and GFP or GFP-Rab3a in melanocyte dendrites. **(D)** Percentage of melanosomes in the vicinity of the plasma membrane [between green and red lines in **(E)]** of melanocyte dendrites. Melanosomes in melanocytes overexpressing GFP or GFP-Rab3a (green) were immunostained (pseudocolored in magenta). Brightfield shows melanosomes as black dots. Arrows indicate co-localization. Scale bars, 10 µm. Plots display mean ± SD of three independent experiments. **(C)** Two-way ANOVA; **(D)** Unpaired t-test. n.s. non-significant; ***P<0.0001. At least 30 images containing approximately 50 melanocytes in total were analyzed per condition.

### Melanin transfer in melanocyte/keratinocyte co-cultures is independent of Rab3a

To further investigate melanin transfer, we tested melanocyte/keratinocyte co-cultures, the system we used in our previous report (Tarafder *et al*., 2014). First, we efficiently silenced Rab3a or Rab11b in melanocytes using siRNA pools (≥75% silencing, Figures S2C and S2D) and then quantified melanin transferred from melanocytes to keratinocytes. As controls, we used both non-transfected melanocytes and melanocytes transfected with a non-targeting siRNA (siControl). We observed that Rab3a depletion does not affect melanin transfer from melanocytes to keratinocytes (Figure 5). Conversely and consistently with our previous studies (Tarafder *et al*., 2014; Moreiras *et al*., 2019), Rab11b silencing impairs melanin transfer by 50% compared with siControl (Figure 5). These results further support the existence of two distinct secretory pathways: one involved in melanocyte/keratinocyte co-cultures controlled by Rab11b, and another stimulated by KCM and regulated by Rab3a.

**Figure 5:**
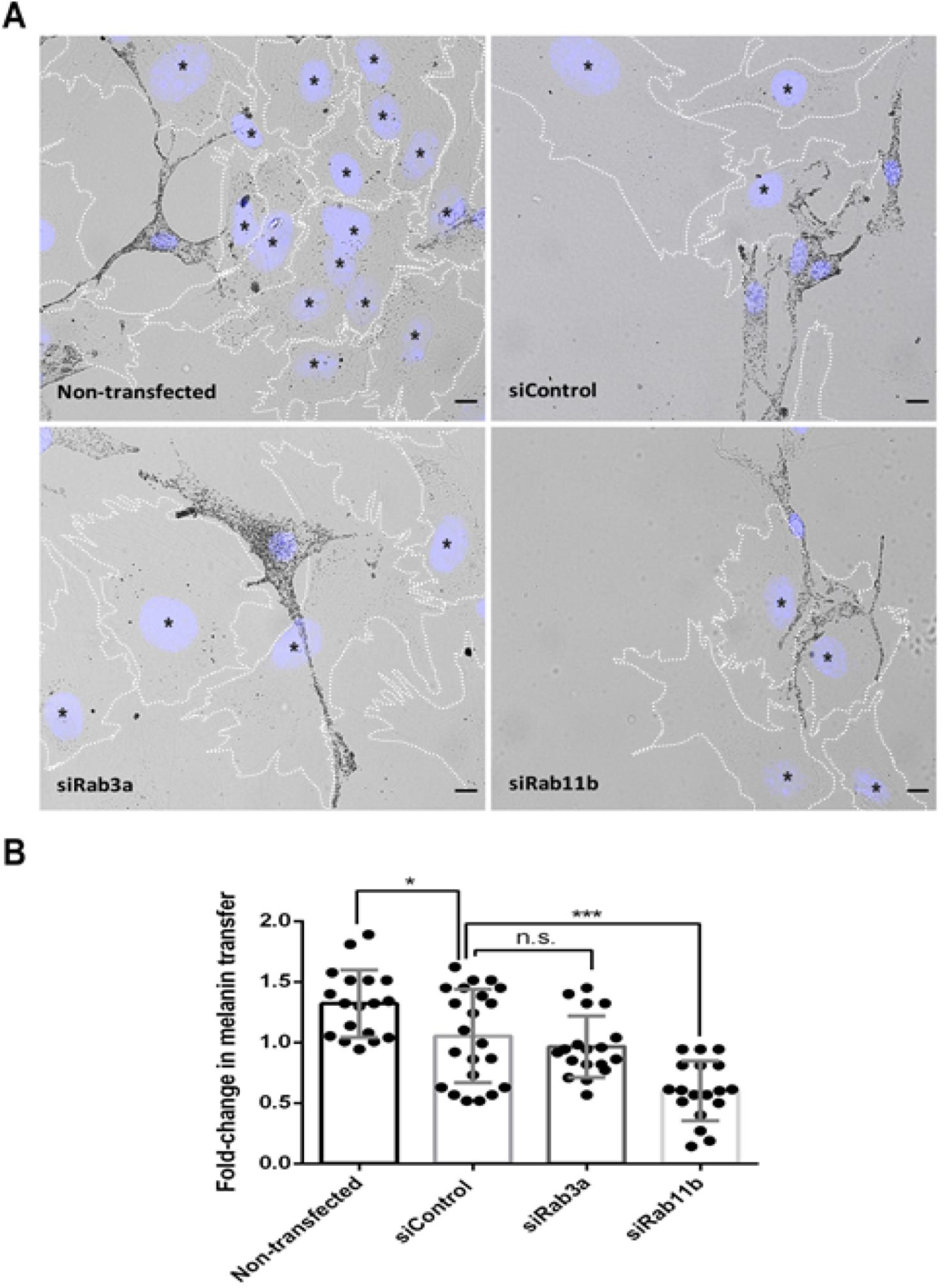
Melanin transfer in melanocyte/keratinocyte co-cultures is Rab11b-dependent and Rab3a-independent. Before performing the co-cultures, melanocytes were silenced for Rab3a (siRab3a) or Rab11b (siRab11b), transfected with a non-targeting siRNA (siControl) or non-transfected. **(A)** Representative brightfield microscopy images of melanocyte/keratinocyte co-cultures with nuclei of keratinocytes labelled with an asterisk (*). **(B)** Fold-change in melanin transfer relative to siControl. At least 60 keratinocytes were analyzed for each condition and the number of melanin-containing organelles per keratinocyte was counted using ImageJ software. Cells were stained with DAPI to label nuclei (blue). White dashed lines define keratinocyte boundaries. Scale bars, 10 μm. The plot represents mean ± SD of three independent experiments. One-way ANOVA (n.s. non-significant; *P<0.05; *** P<0.0001).

## DISCUSSION

Melanin transfer from melanocytes to keratinocytes is a crucial process for skin pigmentation and photoprotection against UVr-induced cell damage. Our previous studies suggested that coupled exo/phagocytosis is the predominant mechanism of melanin transfer in human skin and showed an essential role for Rab11b and the exocyst tethering complex in this process (Tarafder *et al*., 2014; Moreiras *et al*., 2019) ^(1)^. Here we provide evidence that the process of melanin exocytosis is more complex than previously appreciated and demonstrate the existence of at least two types of melanin exocytosis pathways, including a novel one induced by keratinocytes and regulated by the Rab3a small GTPase.

Despite its importance, the molecular mechanisms regulating melanin exocytosis remain poorly characterized, also due to technical limitations and lack of a suitable method to quantify exocytosed melanin. To this end, we adapted and optimized the fluorescence-based method previously published (Fernandes *et al*., 2016) to the measurement of exocytosed melanin from cultured melanocytes with high specificity and sensitivity.

It has been shown that keratinocytes release soluble factors that increase melanin synthesis and transport in melanocytes (Yamaguchi and Hearing, 2010; Taisuke and Hearing, 2011; Hirobe, 2014). However, their effects on melanin exocytosis or transfer from melanocytes to keratinocytes have not been studied (Yamaguchi and Hearing, 2010; Taisuke and Hearing, 2011; Hirobe, 2014). To clarify this unsolved question, we started by assessing the effect on melanin exocytosis of three different culture media: RPMI (Melan-ink4a melanocyte growth medium), DMEM (XB2 keratinocyte growth medium) and KCM (DMEM medium consumed by confluent XB2 keratinocytes in culture for 3 days). We found that soluble factors present in KCM specifically stimulate melanin exocytosis, in addition to stimulating melanogenesis, while DMEM was particularly effective in stimulating melanogenesis and less effective in stimulating melanin exocytosis (Figures 1A and 1B). To explain the effects on melanogenesis, we note that DMEM has 3.5 times more L-tyrosine than RPMI and that L-tyrosine is the initial substrate of tyrosinase, the key melanogenic enzyme (Yamaguchi and Hearing, 2010; Taisuke and Hearing, 2011; Hirobe, 2014). Interestingly, melanocytes cultured with KCM show half of intracellular melanin levels than DMEM (Figures 1B), which can be explained by the increase in melanin exocytosis and/or the consumption of the L-tyrosine in DMEM due to the culture of keratinocytes for 3 days in this medium to prepare KCM. In either case, our results strongly suggest that keratinocyte-derived soluble factors present in KCM stimulate melanin exocytosis from melanocytes in a specific manner that cannot be explained by an increase in melanin synthesis.

Whilst we confirmed that Rab11b regulates melanin exocytosis from melanocytes cultured with RPMI as previously reported by us (Moreiras *et al*., 2019), we found surprisingly that Rab11b does not regulate KCM-stimulated melanin exocytosis (Figures 2A, 2B and 6). Among the candidate Rab GTPases that could regulate melanosome exocytosis, we focused on Rab3a as it localizes to mature melanosome membranes (Araki *et al*., 2000) and is present in melanosome-containing fractions together with several SNAREs involved in exocytic events (Scott and Zhao, 2001). Our studies suggest that Rab3a specifically regulates KCM-induced melanin exocytosis from melanocytes, but not from melanocytes cultured with DMEM or RPMI (Figures 2A, 2B and 6). Intriguingly, the molecular regulators of melanin exocytosis from melanocytes cultured with DMEM remain to be identified as neither Rab11b nor Rab3a show any role in this case. We suggest that other yet uncharacterized melanocytic signaling pathways activating melanin exocytosis do exist and incubation with DMEM should serve as the starting point for future studies.

Additionally, we observed that Rab3a depletion in melanocytes does not affect melanin transfer to keratinocytes in co-cultures. In contrast, Rab11b silencing in melanocytes impairs melanin transfer by 50% (Figure 5), which is consistent with our previous studies (Tarafder *et al*., 2014; Moreiras *et al*., 2019). One note of caution is that the conditioned media experiments tested for melanin exocytosis whilst the co-cultures measured the transfer process, i.e., exocytosis and uptake of melanin by keratinocytes (Figure 5 and 6).

**Figure 6:**
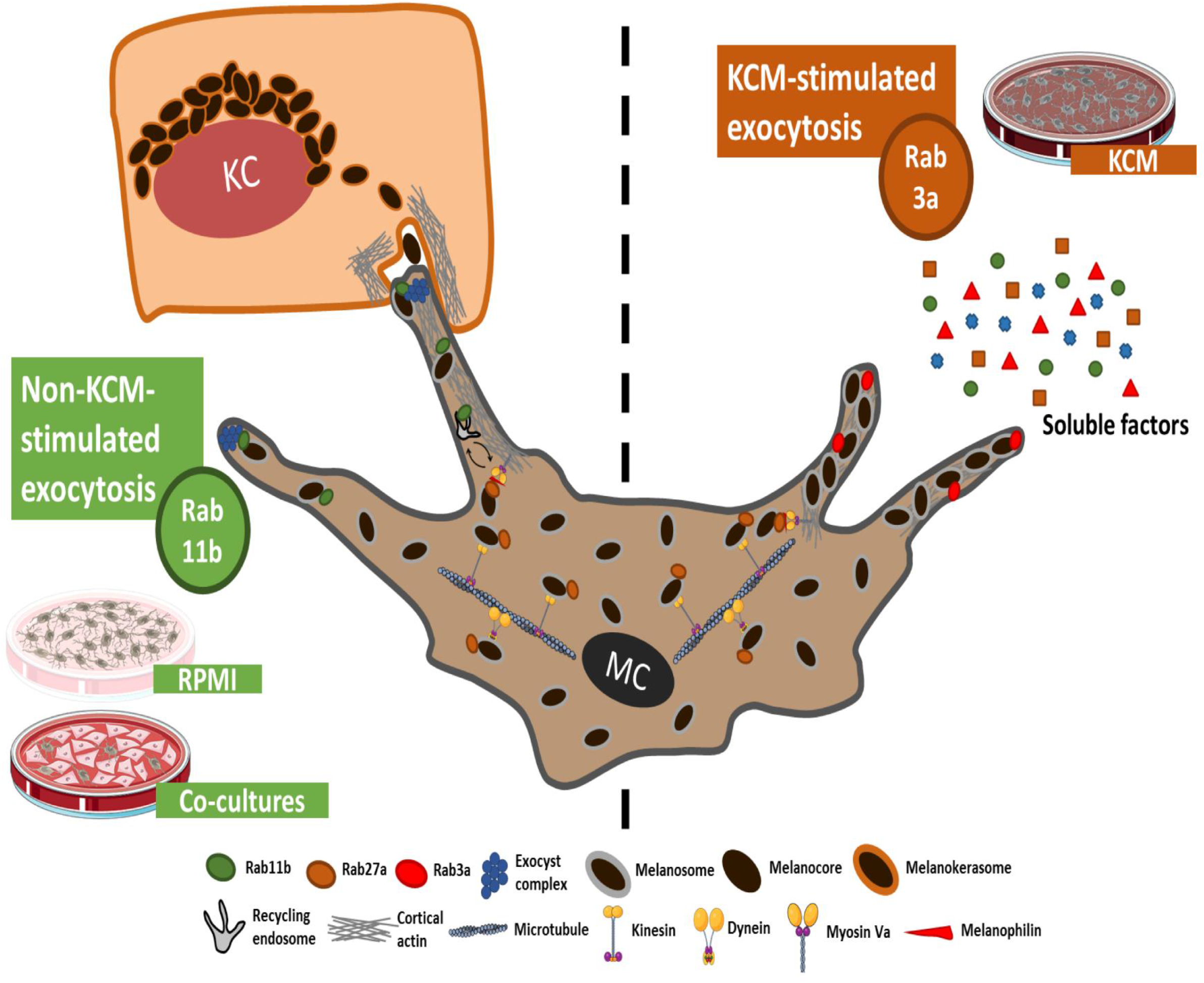
Schematic representation of melanin exocytosis pathways in melanocytes. Melanosomes are tethered to the cortical actin via the tripartite complex Rab27a/Melanophilin/Myosin Va. Melanosomes can then follow two distinct exocytosis pathways, stimulated or not by keratinocyte-conditioned medium (KCM). In the latter (left), Rab11b-positive recycling endosomes are postulated to remodel melanosome membranes, which then interact with the exocyst before being exocytosed as melanocores from dendrites of melanocytes cultured with RPMI, melanocyte/keratinocyte-conditioned medium (MKCM) or co-cultured with keratinocyte. In co-cultures, keratinocytes internalize and store melanocores in specialized organelles, which we proposed to name as melanokerasomes. Alternatively, melanin exocytosis is triggered by KCM (right). This KCM-stimulated route is characterized by a high number of melanosomes in melanocyte dendrites, is Rab3a-dependent and Rab11b-independent.

We also compared melanin exocytosis levels in the presence of keratinocyte-derived factors from MKCM (DMEM medium consumed by confluent Melan-ink4a/XB2 co-cultures for 3 days) or KCM. Melanocytes incubated with MKCM exocytose half the amount of melanin relative to melanocytes cultured with KCM (Figure 1C). However, no significant changes in intracellular melanin levels were observed between these conditions (Figure 1D), reinforcing the specificity of KCM stimulation on melanin exocytosis. Importantly, the crosstalk between melanocytes and keratinocytes in co-cultures can affect the amount or identity of soluble factors secreted by keratinocytes. Indeed, keratinocytes alone and keratinocytes co-cultured with melanocytes show a different secretome. For instance, it was reported that keratinocytes in co-culture release exosomes carrying miRNAs, which in turn stimulate melanogenesis in melanocytes, something that was not observed in keratinocytes cultured alone (Lo Cicero *et al*., 2015). Additionally, co-cultures could allow for cross-talk mediated by cell-to-cell transient contacts. Therefore, the present experiments highlight the importance of melanocyte-keratinocyte communication in the regulation of pigmentation through melanin synthesis, transport and transfer. Moreover, our observations suggest that the concentrations of keratinocyte-derived factors or even the factors themselves are distinct between KCM and MKCM, which can explain the differences in the regulation of melanin exocytosis and exocytosed melanin levels found in each of these conditions (Figure 6). Therefore, it is possible that keratinocytes downregulate the release of soluble factors upon internalizing melanin secreted by melanocytes in co-cultures. In this case, keratinocytes cultured alone would keep secreting factors to stimulate melanin synthesis and exocytosis in a positive feedback loop, resulting in a higher concentration of factors in the medium. These hypotheses should be specifically tested in future studies.

The present results, including the rescue experiments by overexpression of GFP-Rab3a and GFP-Rab3a co-localization with melanosomes close to plasma membrane (Figures 3C and 4C), corroborate the specific role of Rab3a in KCM-stimulated melanin exocytosis. The enrichment of melanosomes in dendrites induced by KCM was also a significant novel finding in this study (Figures 3C and 6). The underlying mechanisms remain unclear but our results suggest that melanocytes are able to coordinately regulate melanogenesis, melanosome peripheral transport and regulated exocytosis. One such mechanism linking transport and exocytosis could be a cooperative role in the dendritic actin-mediated transport by Rab27a (Barral and Seabra, 2004; Hume *et al*., 2007; Wu *et al*., 2012) and exocytosis of melanosomes by Rab3a. Such functional cooperation has been described for other LROs, including Weibel-Palade bodies and dense core granules present in endothelial and sperm cells, respectively (Raposo, G. *et al*., 2007; Zografou *et al*., 2012; Quevedo *et al*., 2018; Delevoye *et al*., 2019).

In summary, our data suggests a complex cross-talk between melanocytes and keratinocytes in the regulation of skin pigmentation. We demonstrate that keratinocyte soluble factors present in KCM activate melanocytic signaling pathways leading to enhanced melanogenesis, peripheral transport and regulated exocytosis. The latter is regulated by Rab3a and not Rab11b, which demonstrates the existence of at least two distinct melanin exocytosis pathways: one controlled by Rab11b, and another stimulated by KCM and regulated by Rab3a (Figure 6). Rab11b regulates melanin exocytosis both in unstimulated melanocytes cultured with RPMI, as well as melanin transfer in melanocyte-keratinocyte co-cultures containing MKCM (Figure 6). The stimulated pathway is specific for melanocytes cultured with KCM, leading to the increase of exocytosed melanin, in addition to stimulating melanogenesis (Figure 6). Future studies should be directed at characterizing the differences between the keratinocyte-derived soluble factors present in KCM and MKCM and identifying the signaling pathway(s) and downstream Rab3a-effectors that lead to the stimulation of KCM-induced melanin exocytosis.

## MATERIALS AND METHODS

### Cell culture

Melan-ink4a mouse melanocytes were cultured in RPMI-1640 (Gibco, Grand Island, New York) supplemented with 10% fetal bovine serum (Gibco, Grand Island, New York), 200 pM cholera toxin (Gentaur, Kampenhout, Belgium), 200 nM phorbol-myristate-acetate (Alfa Aesar, Heysham, England), 100 units/ml penicillin (Gibco, Grand Island, New York) and 100 µg/ml streptomycin (Gibco, Grand Island, New York), at 37°C with 10% CO_2_. XB2 mouse keratinocytes were cultured in DMEM (Gibco, Grand Island, New York) supplemented with 10% fetal bovine serum, GlutaMAX (2 mM L-alanyl-L-glutamine dipeptide - Gibco, Grand Island, New York), 100 units/ml penicillin and 100 µg/ml streptomycin, at 37°C with 10% CO_2_. Melan-ink4a/XB2 co-cultures were incubated in DMEM supplemented with 10% fetal bovine serum, GlutaMAX, 200 pM cholera toxin, 200 nM phorbol-myristate-acetate, 100 units/ml penicillin and 100 µg/ml streptomycin, at 37°C with 10% CO_2_. Both Melan-ink4a and XB2 cells were obtained from the lab of Dorothy Bennett and Elena Sviderskaya (St. George’s Hospital, London) in June 2019 and upon arrival were immediately used. These cell lines used were tested for mycoplasma using a PCR assay performed by the certified entity Eurofins Genomics Europe in July 2020.

### Conditioned media production

To produce keratinocyte-conditioned medium (KCM), 10 ml of complete medium were added to XB2 keratinocytes at 80% confluency and incubated for 3 days. In the case of melanocyte/keratinocyte-conditioned medium (MKCM) production, Melan-ink4a melanocytes and XB2 keratinocytes were seeded in 12-well plates, in DMEM without cholera toxin or phorbol-myristate-acetate at a ratio of 1 melanocyte to 5 keratinocytes. Finally, the co-cultures were incubated until they reached 80% confluency and at that point, the culture medium was replaced by 2 ml of fresh DMEM medium and incubated for further 3 days. After collection, all conditioned media were filtered using a 0.45 μm syringe filter (Sarstedt, Nümbrecht, Germany) and stored at -80°C until they were used. MKCM was centrifuged at 21,000 x *g* for 1.5 hours at 4°C to discard exocytosed melanin, before being stored.

### siRNA transfection

Melan-ink4a melanocytes were cultured at 8.5×10^4^ cells/well in 12-well plates. While 50 nM of SMART pool siRNAs (Thermo Scientific, Lenexa, Kansas, Table S1) were added to 250 μl of Opti-MEM (Gibco, Grand Island, New York), 2.5 μl of DharmaFECT 4 (Dharmacon, Lafayette, Colorado) were added to 250 μl of Opti-MEM. After 5 minutes of incubation at room temperature, the two mixtures were combined, mixed gently, and incubated for 20 minutes at the same temperature. The growth medium from cells seeded the day before transfection was removed and replaced by the siRNA mixture in a total volume of 500 μl, in Opti-MEM. Cells were incubated for 24 hours at 37°C and then the medium was changed to 1 ml of complete RPMI, DMEM or KCM supplemented with 200 pM cholera toxin and 200 nM phorbol-myristate-acetate. After 3 days, cells were assayed and silencing efficiency was confirmed.

### Plasmid transfection

Melan-ink4a melanocytes were seeded on glass coverslips in 24-well plates at 3×10^4^ cells/well and the day after were transfected with 1 μg of cDNA plasmid (Table S2) mixed with 2 μl of FuGENE6 (Promega, Madison, Wisconsin) in 500 μl Opti-MEM. Cells were incubated for 4 hours at 37°C and then the medium was changed to 1 ml DMEM or KCM supplemented with 200 pM cholera toxin and 200 nM phorbol-myristate-acetate. After 1 day, cells were analyzed by confocal microscopy.

### Lentiviral production and cell transduction

The lentiviral vector pLenti6/V5-DEST Gateway was used to achieve stable overexpression of GFP or GFP-Rab3a. For this, the cDNA sequences were inserted in the lentiviral vector. For lentivirus production, STAR-RDpro cells grown in DMEM supplemented with 10% FBS, 2 mM GlutaMAX and 15 mM HEPES were co-transfected using jetPRIME® (Poly Plus, Berkeley, California) with 2.5 µg of lentiviral vector, 3.5 µg packaging plasmid (pxPAX2) and 1.8 µg envelope plasmid (pCMV-VSV-G). Media containing lentiviral particles were harvested 48 hours post-transfection and stored in aliquots at -80°C until use. Melan-ink4a cells were plated at 8.5 × 10^4^ cells/well in 12-well plates and transduced on the next day with approximately 3 × 10^5^ plaque-forming unit (PFU) of lentiviral particles in the presence of 8 µg/ml polybrene (hexadimethrine bromide, Sigma-Aldrich, Steinheim, Germany). After 24 hours, 9 µg/ml blasticidin were added to transduced cells. Cells were selected for at least 5 days before assayed.

### qRT-PCR quantification of gene expression

Total RNA was isolated from cells using the RNeasy Mini kit (Qiagen, Venlo, Netherlands) and converted to cDNA using SuperScript® II (Invitrogen, Lenexa, Kansas) according to the manufacturers’ instructions. Real-time quantitative PCR (qRT-PCR) reactions were performed using a Roche LightCycler (Roche, Grenzacherstrasse, Switzerland) and Roche SybrGreen Master Mix reagent (SybrGreen, Roche, Grenzacherstrasse, Switzerland). For each gene, specific primers were used (Table S3). Gene expression was calculated relative to control cells and normalized to *Gapdh*.

### Melanin quantification

Wild-type Melan-ink4a melanocytes or lentivirus-transduced Melan-ink4a melanocytes overexpressing GFP or GFP-Rab3a were seeded in 12-well plates at 8.5×10^4^ cells/well. On the next day, melanocytes were silenced using SMART pool siRNAs (Table S1). Then, cells were incubated for 3 days with RPMI, DMEM or KCM supplemented with 200 pM cholera toxin and 200 nM phorbol-myristate-acetate. The culture medium was collected and centrifuged at 300 x *g* for 5 minutes to remove dead cells. The supernatant was collected and centrifuged at 21,000 x *g* at 4 °C for 1.5 hours to precipitate melanin. Afterwards, the pellet was resuspended in 90 μl of NaOH 1 M (LabKem, Migjorn, Barcelona) with 10% dimethyl sulfoxide (DMSO, Sigma-Aldrich, Steinheim, Germany).

To quantify intracellular melanin, cells were trypsinized and collected by centrifugation at 300 x *g* for 5 minutes. The pellet obtained was washed with phosphate-buffered saline (PBS), centrifuged again at 300 x *g* for 5 minutes and then lysed in 500 μL of NaOH 1 M + 10% DMSO. Melanin samples, including synthetic melanin (Sigma-Aldrich, Steinheim, Germany) to obtain the standard curve were solubilized in NaOH 1M + 10% DMSO, boiled at 80°C for 1 hour and centrifuged at 3,000 x *g* for 5 minutes to remove insoluble debris. Then, 50% (w/v) H_2_O_2_ (Acros Organics, Lenexa, Kansas) was added to all the samples to a final concentration of 30% (v/v) and incubated for 4 hours at room temperature in the dark. Finally, 200 μL of sample were added to black opaque 96-well plates (Greiner, Kremsmünster, Austria) and read in a SpectraMax i3x multi-mode microplate reader (FortéBio, Goettingen, Germany) using an optimized reading distance of 2.81 mm and excitation/emission wavelengths of 470 nm/550 nm. The amount of melanin for each sample was normalized to the total protein. For this, the protein content in the cell lysates was measured at 562 nm in the same plate reader, using the Pierce BCA protein kit (Thermo Scientific, Lenexa, Kansas) and following the manufacturer’s protocol. Exocytosed and intracellular melanin amounts were calculated by interpolating their fluorescence values with the ones from the standard curve of synthetic melanin, considering NaOH 1M + 10% DMSO as blank and normalizing to the protein levels of each sample.

### Melanin transfer assay

Melan-ink4a melanocytes were seeded in 12-well plates and transfected with SMART pool siRNAs (Table S1). On the following day, Melan-ink4a were detached and re-seeded (6.67×10^3^ cells/well) with XB2 keratinocytes (3.33×10^4^ cells/well) at a ratio of 1:5 in 24-well plates containing glass coverslips. After 3 days, the coverslips were washed 3 times with PBS and fixed with 4% paraformaldehyde (PFA, Alfa Aesar, Heysham, England) in PBS for 20 minutes at room temperature. Finally, the coverslips were mounted with Fluoromount-G with DAPI (Invitrogen, Lenexa, Kansas) to visualize the cells’ nuclei and images were acquired in a ZEISS Axio Imager 2 microscope with a PlanApochromat 63×1.4 NA oil-immersion objective. The melanin-containing organelles present within keratinocytes, which we proposed to call melanokerasomes (Moreiras *et al*., 2021b), were counted using ImageJ software and normalized to the number of cells analyzed in each image.

### Immunofluorescence microscopy

Melan-ink4a melanocytes were seeded on glass coverslips in 24-well plates (3×10^4^ cells/well) and transfected with different plasmids (Table S2). On the next day, the coverslips were washed 3 times with PBS and cells fixed with 4% paraformaldehyde (PFA, Alfa Aesar, Heysham, England) in PBS for 20 minutes. Blocking and permeabilization were done by incubating the cells for 30 minutes with Perblock solution (0.5% BSA and 0.1% Saponin in PBS). Coverslips were then incubated with the primary antibodies anti-HMB45 (Dako M0634, Santa Clara, California) or anti-COPI (hybridoma clone M3A5), diluted 1:200 in Perblock solution for 1.5 hours in a humidified chamber. After washing 3 times with PBS, coverslips were incubated with Alexa 568-conjugated goat anti-mouse secondary antibody (Invitrogen, Lenexa, Kansas) diluted 1:500 in Perblock solution for 45 minutes in a humidified chamber. Finally, coverslips were washed 3 times with PBS and mounted with Fluoromount-G with DAPI (Invitrogen, Lenexa, Kansas) to visualize the cells’ nuclei. All the steps were performed at room temperature. The images were acquired in a Zeiss LSM 710 confocal microscope with a PlanApochromat 63×1.4 NA oil-immersion objective and analyzed with ImageJ software or Icy 2.0.2.0 software. For co-localization analysis, all microscopy images were pre-processed with a mask using the threshold tool in ImageJ software and Mander’s overlap coeficient was calculated using the plugin Coloc 2 of the same software. The Icy 2.0.2.0 software was used to calculate the percentage of melanosomes in three distinct regions of interest of melanocytes (cell body, dendrites and regions in the vicinity of plasma membrane in dendrites). For this, the regions were outlined with free form lines and the number of melanosomes in each one was counted using the Spot detector tool. In order to normalize the region close the plasma membrane between distinct images, its thickness was defined by reducing automatically in 30% the area outlining dendrites using the rescale tool of the software. Finally, to calculate the percentage of melanosomes, the number in each of three regions was normalized to the number of melanosomes in the whole cell.

### Immunoblotting

Melan-ink4a melanocytes plated on 12-well plates and transfected/transduced with siRNAs and/or lentiviruses, were lysed in ice-cold lysis buffer (50 mM Tris-HCl, pH 7.5, 150 mM NaCl, 1 mM EDTA, 1 mM EGTA, 2 mM MgCl2, 1 mM DTT and 1% IGEPAL) in the presence of protease inhibitors for 30 minutes on ice, followed by centrifugation at 21,000 x *g* for 30 minutes at 4 °C. Total protein concentration was quantified using the DC protein assay kit (Bio-Rad, Hercules, California) following the manufacturer’s protocol, and equal protein amounts per sample were resuspended in 2× loading buffer. Then, samples were resolved on 12% SDS-PAGE and the protein bands transferred onto activated nitrocellulose membranes at 100 V in transfer buffer (25 mM Tris, 192 mM glycine, and 20% ethanol) for 55 minutes at room temperature. Membranes were blocked with blocking buffer (5% non-fat dry milk and 0.1% Tween-20 in PBS) and incubated with the primary polyclonal antibodies goat anti-Rab3a (AB0032-100, SICGEN, Lisbon, Portugal) diluted 1:1,000 in PBS andor goat anti-Gapdh (AB0049-500, SICGEN, Lisbon, Portugal) diluted 1:1,000 in blocking buffer. Both primary antibodies were separately incubated overnight at 4°C under constant mixing. After each incubation, membranes were washed 3 times with PBS + 0,1% Tween-20 and incubated, for 1 hour at room temperature, with HRP-conjugated secondary anti-goat antibody (A16005, Thermo Scientific, Lenexa, Kansas) diluted 1:7,000 in blocking buffer. Antibody complexes were visualized by employing the enhanced chemiluminescence reagent (ECL, Amersham Biosciences, Waukesha, Wisconsin). The signal was detected using an Imager Chemidoc XRS (Bio-Rad, Hercules, California) and the images were pre-processed with Image Lab software (Bio-Rad, Hercules, California). Band intensities were quantified using ImageJ software by generating peaky histograms that are proportional to area and intensity of bands. The housekeeping protein Gapdh was used to normalize Rab3a levels and the results indicated as percentage relative to control. Finally, membrane stripping was done by incubating twice with a mild stripping buffer (1.5% glycine, 0.1% SDS and 1% Tween, pH 2.2) for 5 minutes at room temperature. Stripped membranes were then incubated with primary polyclonal antibody goat anti-GFP (AB0020-500, SICGEN, Lisbon, Portugal), diluted 1:1,000 in blocking buffer, followed by HRP-conjugated secondary anti-goat, and developed as described above.

### Statistical analysis

All numerical data are representative of three or more biological replicates and presented as mean ± standard deviation (SD). One-way ANOVA (Turkey’
ss multiple comparisons test) or two-way ANOVA (Turkey’s multiple comparisons test) were applied to data with one independent variable or two independent variables, respectively. Where indicated, one-way ANOVA (Dunnett’s multiple comparison test) was used to compare different data sets with control. Unpaired t-test with Welch’s correction was applied to the plots showing two conditions with only one independent variable. The statistical analyses were performed using GraphPad Prism software version 6.01.

## DATA AVAILABILITY STATEMENT

No datasets were generated or analyzed during the current study.

## ACKNOWLEDGEMENTS

We would like to thank our group for the critical reading of the manuscript and CEDOC microscopy and cell culture facilities. This study was supported by Fundação para a Ciência e a Tecnologia (FCT), Portugal through grant PTDC/BIA-CEL/29765/2017 and PhD fellowships to LCC, MVN and LBL (DFA/BD/8812/2020, PD/BD/137442/2018 and SFRH/BD/131938/2017, respectively). This work was developed with the support from the research infrastructure PPBI-POCI-01-0145-FEDER-022122, co-financed by FCT (Portugal) and Lisboa2020, under the PORTUGAL2020 agreement (European Regional Development Fund). This article is supported by the LYSOCIL project, which has received funding from the European Union’s Horizon 2020 research and innovation programme under grant agreement No. 811087.

## AUTHOR CONTRIBUTIONS STATEMENT

LCC - conceptualization, formal analysis, investigation, methodology, writing (original draft) and editing; LBL - conceptualization, investigation, methodology and writing (original draft); MVN - investigation, methodology and writing (original draft); JSR - resources and methodology; MCS - conceptualization, supervision, resources, investigation, methodology and writing (original draft); DCB - conceptualization, formal analysis, supervision, funding acquisition, resources, investigation, methodology, project administration, writing (original draft) and editing.

## Footnotes

1 Moreiras, H., Neto, M. V, Bento-Lopes, L., Escrevente, C., Ramalho, J. S., Seabra, M. C. & Barral, D. C. (2021a). Melanocore uptake by keratinocytes occurs through phagocytosis and involves Protease-activated receptor-2 activation. bioRxiv, 2021.04.13.439501. https://doi.org/10.1101/2021.04.13.439501.

