## Supplementary Figures and Tables for "Rab3a regulates melanin exocytosis induced by keratinocyte-conditioned medium"

### Figure S1

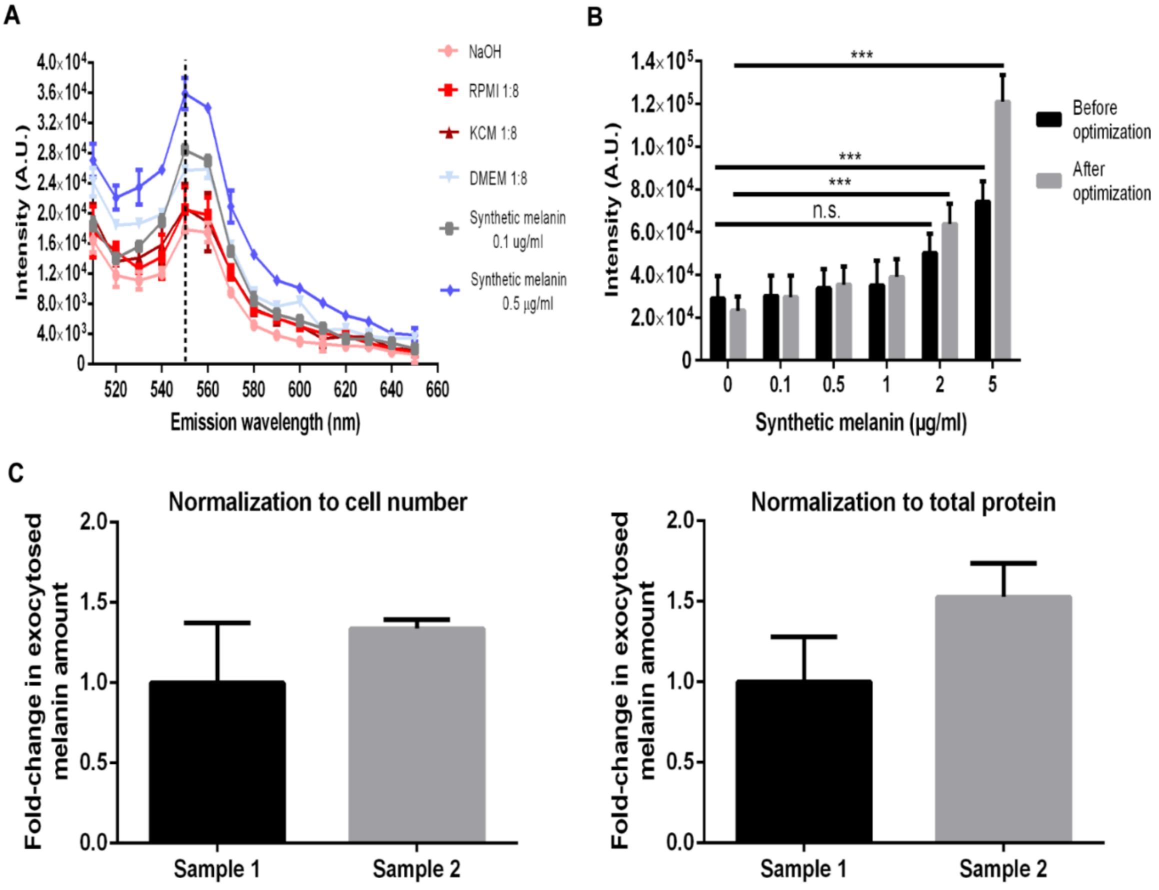


***Figure S1: Optimization of a specific and sensitive method to quantify exocytosed melanin.*** **(A)** Comparison between the emission spectra of culture media (RPMI, DMEM and KCM) diluted 1:8 in NaOH and the emission spectrum of 0.1 μg/ml or 0.5 μg/ml of synthetic melanin. **(B)** Comparison between synthetic melanin standards and indicated concentrations before and after optimizing the final sample volume and the reading distance. **(C)** Exocytosed melanin levels quantified in two samples (Sample 1 and 2) and normalized to the number of cells or the total protein in each sample. Results are represented in arbitrary units (A.U.) or fold-change. The plots represent mean ± SD of 3 independent experiments. Two-way ANOVA (n.s. non-significant; *** $P<0.0001$).

**Figure S2**

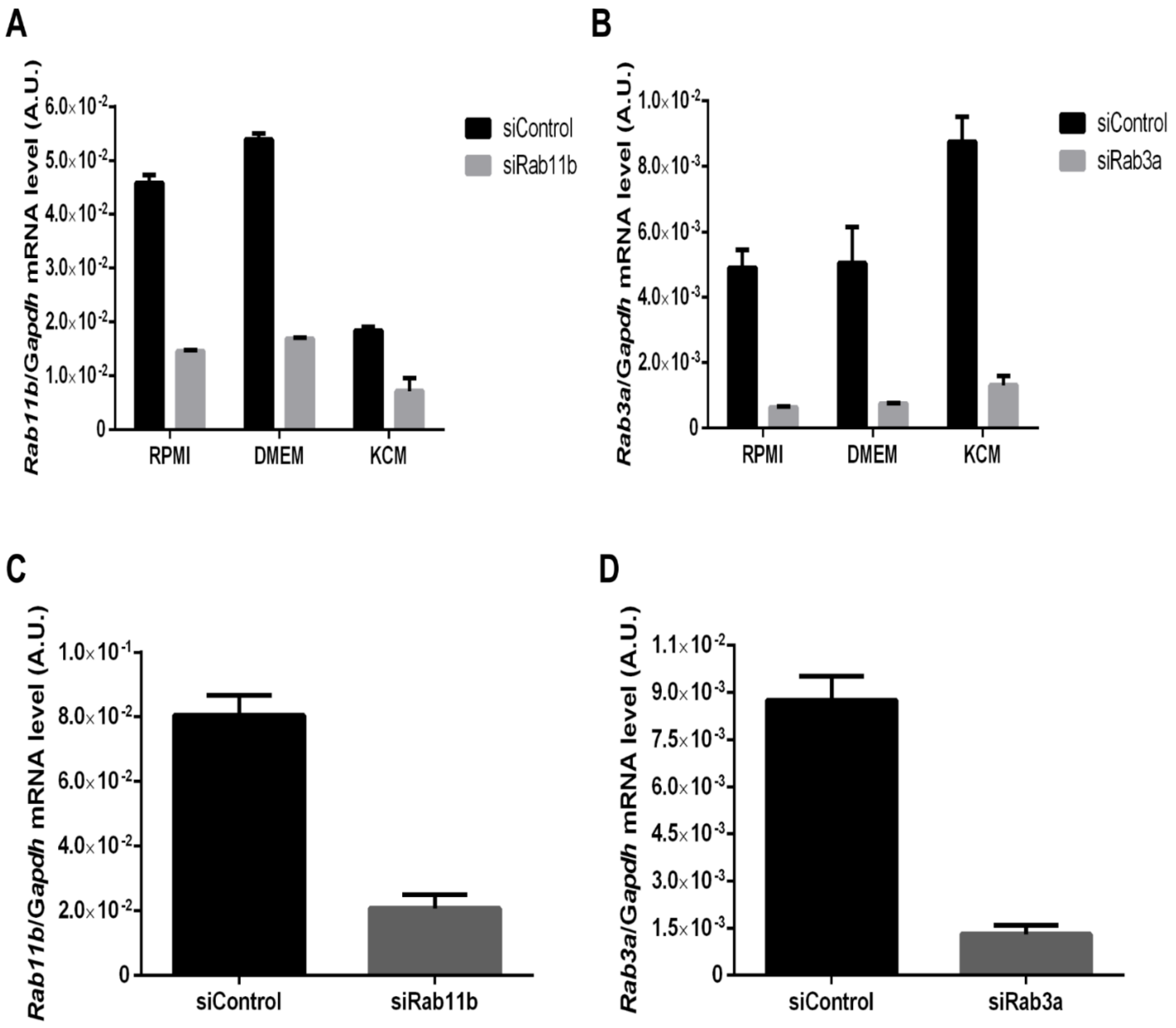


***Figure S2: Silencing efficiency of Rab11b and Rab3a in Melan-ink4a melanocytes.*** **(A)** Rab11b or **(B)** Rab3a silencing in melanocytes cultured with RPMI, DMEM or KCM. **(C)** Rab11b or **(D)** Rab3a silencing in melanocytes used to perform melanocyte/keratinocyte co-cultures. Plots show values of relative gene expression normalized to *Gapdh*. A.U., arbitrary units. Error bars represent mean ± SD of three independent experiments.

**Figure S3**

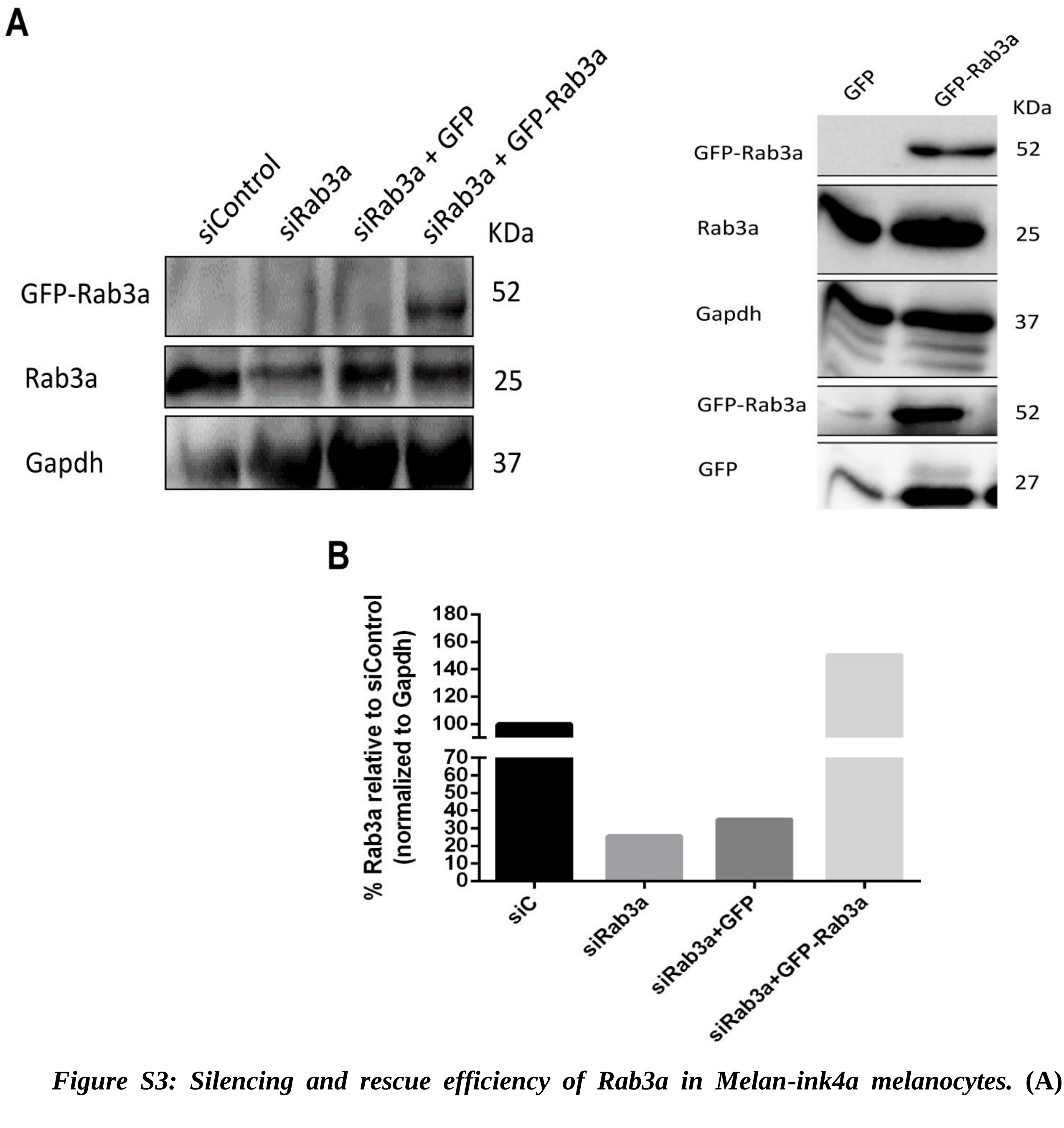


***Figure S3: Silencing and rescue efficiency of Rab3a in Melan-ink4a melanocytes.*** **(A)** Representative western blot for GFP-Rab3a, GFP and Rab3a. Gapdh was used as a loading control. **(B)** Quantification of total Rab3a protein levels normalized to Gapdh and shown in percentage relative to the control.

**Figure S4**

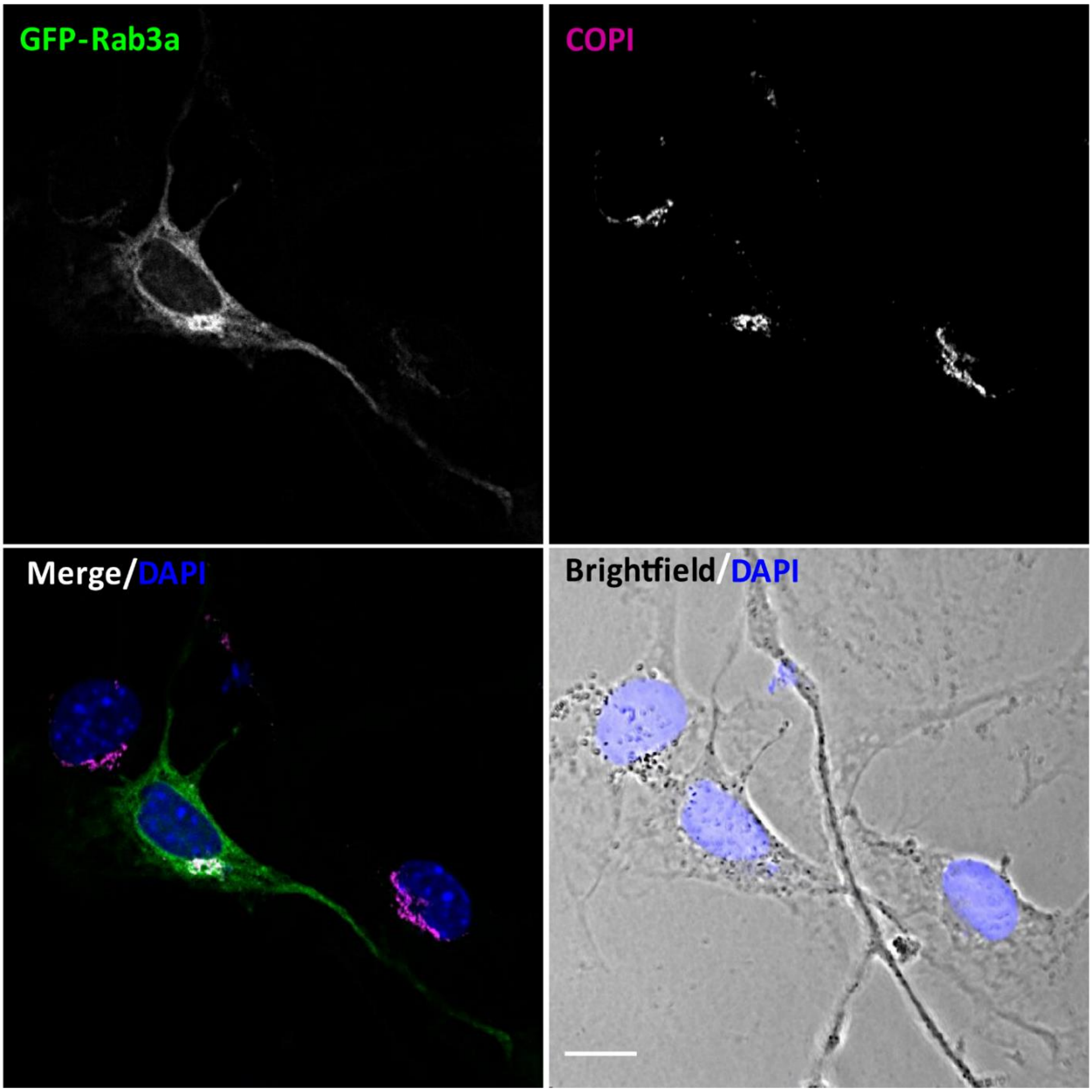


***Figure S4: Rab3a accumulates at the Golgi in the perinuclear region***. Representative confocal images of Melan-ink4a melanocytes overexpressing GFP-Rab3a (green) and immunostained for the Golgi marker COPI (pseudocolored in magenta). Images were analyzed with ImageJ software. Melanocytes were stained with DAPI to label nuclei (blue) and the brightfield shows melanosomes as black dots. Scale bar, 10 µm. Images are representative of three independent experiments.

**<u>Table S1</u>** - ***List of SMART pool siRNAs used.***

| **siRNAs** | **Species** | **Sequences (5'-3')** |
|---|---|---|
| Rab3a | Mouse | CCACUCAGAUCAAAACUUA<br>UGUCAGCACCGUUGGCAUA<br>GGACGUGAUCUGUGAGAAG<br>CCAUCUACCGCAACGACAA |
| Rab11b | Mouse | ACAGAAAUCUACCGUAUUG<br>CGAGUACGAUUACCUAUUC<br>GCAGAUAGCAACAUUGUCA<br>GUGCACUGCUGGUAUAUGA |
| Control | Mouse | GAAGAUAUCGUCCGCAUUA<br>GUUGAAAUUUGACCAGUUA<br>GAACUUAGCAGGAGAGAGU<br>CUUCAGAUGUGUUCAAGAA |

**<u>Table S2</u>** – ***List of cDNA plasmids used.***

| | **Plasmids** |
|---|---|
| **GFP** | pENTR GFP vector |
| **GFP-Rab3a** | pENTR GFP C2 mRab3a |

**<u>Table S3</u>** – ***List of primers used.***

| **Primers** | **Species** | **Sequences (5'-3')** |
|---|---|---|
| Rab3a Forward<br>Rab3a Reverse | Mouse | TTAATGCAGTGCAGGACTGG<br>CAGGCACAATCCTGATGAGG |
| Rab11b Forward<br>Rab11b Reverse | Mouse | AAGGAGCTGCGGGATCATGC<br>ACAGGCTCTGGCAGCACTGC |
| Gapdh Forward<br>Gapdh Reverse | Mouse | AACTTTGGCATTGTGGAAG<br>ACACATTGGGGGTAGGAAC |
